# The effect of sample size on polygenic hazard models for prostate cancer

**DOI:** 10.1101/679092

**Authors:** Roshan A. Karunamuni, Minh-Phuong Huynh-Le, Chun C. Fan, Rosalind A. Eeles, Douglas F. Easton, ZSofia Kote-Jarai, Ali Amin Al Olama, Sara Benlloch Garcia, Kenneth Muir, Henrik Gronberg, Fredrik Wiklund, Markus Aly, Johanna Schleutker, Csilla Sipeky, Teuvo LJ Tammela, Børge G. Nordestgaard, Tim J. Key, Ruth C. Travis, David E. Neal, Jenny L. Donovan, Freddie C. Hamdy, Paul Pharoah, Nora Pashayan, Kay-Tee Khaw, Stephen N. Thibodeau, Shannon K. McDonnell, Daniel J. Schaid, Christiane Maier, Walther Vogel, Manuel Luedeke, Kathleen Herkommer, Adam S. Kibel, Cezary Cybulski, Dominika Wokolorczyk, Wojciech Kluzniak, Lisa Cannon-Albright, Hermann Brenner, Ben Schöttker, Bernd Holleczek, Jong Y. Park, Thomas A. Sellers, Hui-Yi Lin, Chavdar Slavov, Radka Kaneva, Vanio Mitev, Jyotsna Batra, Judith A. Clements, Amanda Spurdle, Australian Prostate Cancer BioResource (APCB), Manuel R. Teixeira, Paula Paulo, Sofia Maia, Hardev Pandha, Agnieszka Michael, Ian G. Mills, Ole A. Andreassen, Anders M. Dale, Tyler M. Seibert, The PRACTICAL Consortium

## Abstract

We aimed to determine the effect of sample size on performance of polygenic hazard score (PHS) models in predicting the age at onset of prostate cancer. Age and genotypes were obtained for 40,861 men from the PRACTICAL consortium. The dataset included 201,590 SNPs per subject, and was split into training (34,444 samples) and testing (6,417 samples) sets. Two PHS model-building strategies were investigated. Established-SNP model considered 65 SNPs that had been associated with prostate cancer in the literature. A stepwise SNP selection was used to develop Discovery-SNP models. The performance of each PHS model was calculated for random sizes of the training set (1 to 30 thousand). The performance of a representative Established-SNP model was estimated for random sizes of the testing set (0.5 to 6 thousand). Mean HR_98/50_ (hazard ratio of top 2% to the average in the test set) of the Established-SNP model increased from 1.73[95%CI: 1.69-1.77] to 2.41[2.40-2.43] when the number of training samples was increased from 1 to 30 thousand. The corresponding HR_98/50_ of the Discovery-SNP model increased from 1.05[0.93-1.18] to 2.19[2.16-2.23]. HR_98/50_ of a representative Established-SNP model using testing set sample sizes of 0.6 and 6 thousand observations were 1.78[1.70-1.85] and 1.73[1.71-1.76], respectively. We estimate that a study population of 20 to 30 thousand men is required to develop Discovery-SNP PHS models for prostate cancer. The required sample size could be reduced to 10 thousand samples, if a set of SNPs associated with the disease has already been established.

**Author summary:** Polygenic hazard scores represent a recent advancement in polygenic prediction to model the age of onset of various diseases, such as Alzheimer’s disease or prostate cancer. These scores accumulate small effect sizes from several tens of genetic variants and can be used to establish an individual’s risk of experiencing an event relative to a control population across time. The largest barrier to the development of polygenic hazard scores is the large number of study subjects needed to develop the underlying models. We sought to understand the effect of varying the total number of samples on the performance of a polygenic hazard score in the context of prostate cancer. We found that the performance of the score did not appreciably change beyond 20 to 30 thousand observations when developing the model from scratch. However, when the discovery of the genetic variants can be borrowed from those already identified in the literature to be associated with the disease, the required number of samples is reduced to 10 thousand with no appreciable detriment in performance. We hope that these results can guide the design of future studies of polygenic scores in other diseases and demonstrate the importance of genome-wide association studies.

## Introduction

Polygenic prediction models have been studied extensively for several diseases such as prostate cancer[1], breast cancer[2], type 2 diabetes[3], dementia[4], and atherosclerosis[5]. Polygenic scores in the context of survival models are a more recent advancement in the field, but have been garnering interest in the prediction of age at onset of Alzheimer’s disease[6] and prostate cancer[7]. The steady increase in genetic testing[8,9], both in public and clinical domains, suggests that survival models could be applied to new diseases. The largest obstacle to the development of these models is the large number of study subjects, often in the tens of thousands[8], which are required for robust training and testing.

Our aim was to quantify the effect of sample size on the performance of a polygenic survival model. This was explored through a specific disease condition that is expected to be representative, namely the prediction of age of onset in prostate cancer. We investigated two potential model development strategies. For the ‘Established-SNP’ model, we selected single-nucleotide polymorphisms (SNPs) that had previously been shown to be associated with prostate cancer, and simply estimated the coefficients for these SNPs in a Cox proportional hazards framework. For the ‘Discovery-SNP’ model, we implemented the SNP selection technique described by Seibert *et al.*[7] to identify SNPs in our genotyping data for inclusion in the Cox proportional hazards framework. The Established-SNP and Discovery-SNP represent two strategies that researchers could employ to build a polygenic survival model. In order to simulate samples of different sizes, we randomly sampled our training and testing sets. The results of this work will help inform the design of future studies to develop polygenic survival models for other diseases.

## Results

### Established- vs. Discovery-SNP model performance

Histogram comparisons of performance metrics of Established (EST) and Discovery (DIS) SNP models are illustrated in Figure 1. The performance metrics are shown for 50 random samplings of the training set using a sample size of 30 thousand total observations. Qualitatively, there appears to be more variability in performance metrics associated with the Discovery process.

**Figure 1.**
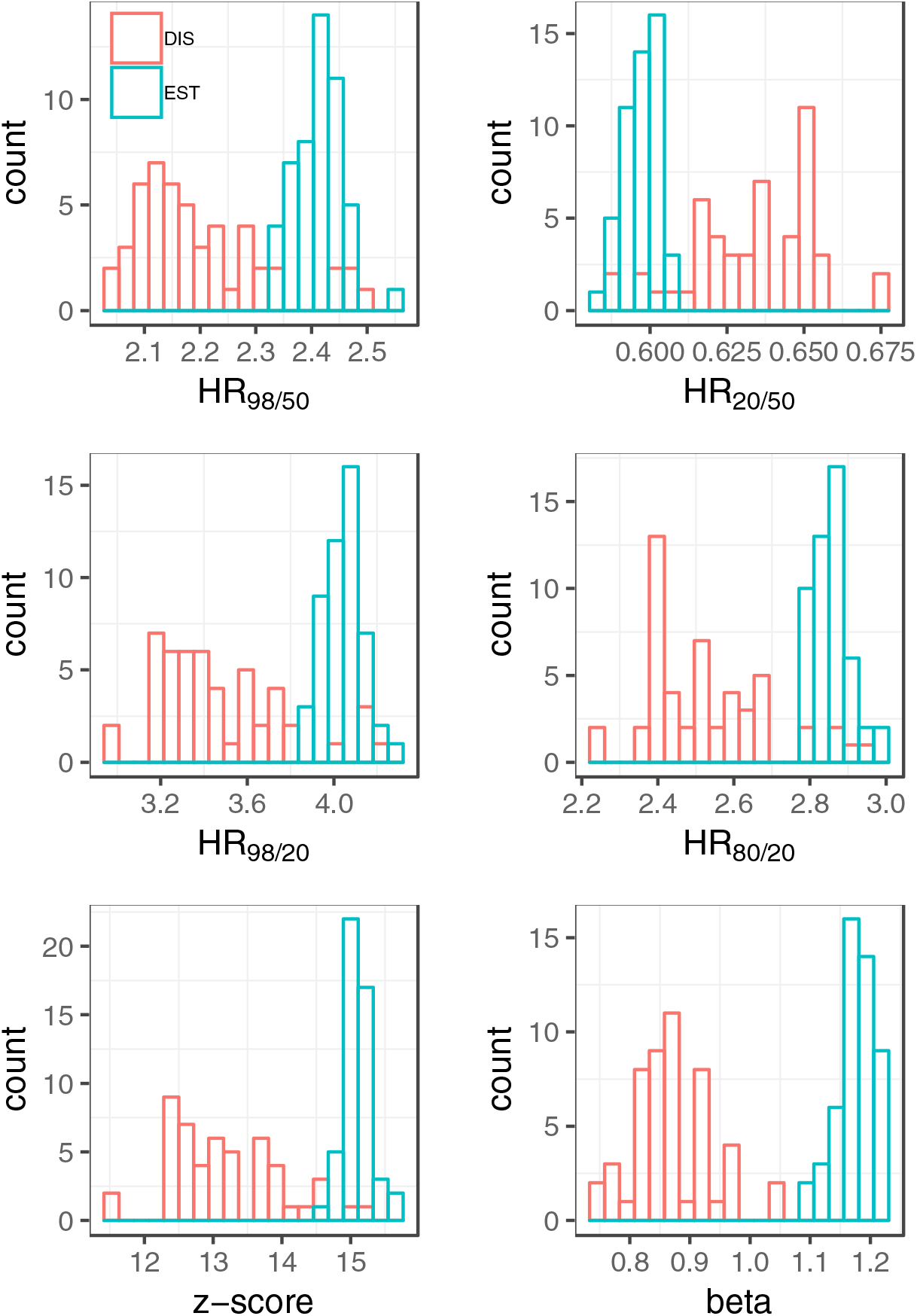
Comparison of performance metrics between Established (EST) and Discovery (DIS) SNP models using 50 random samples of the training set using a sample size of 30 thousand. There is more variability with the Discovery process. Established SNPs, though, were discovered using the data in the training set; this circularity is not accounted for in the present study, which focuses on sample size effects.

### Coefficients of Established-SNP model

The mean coefficients for the 65 SNPs used in the Established-SNP model are plotted in Figure 2.

**Figure 2.**
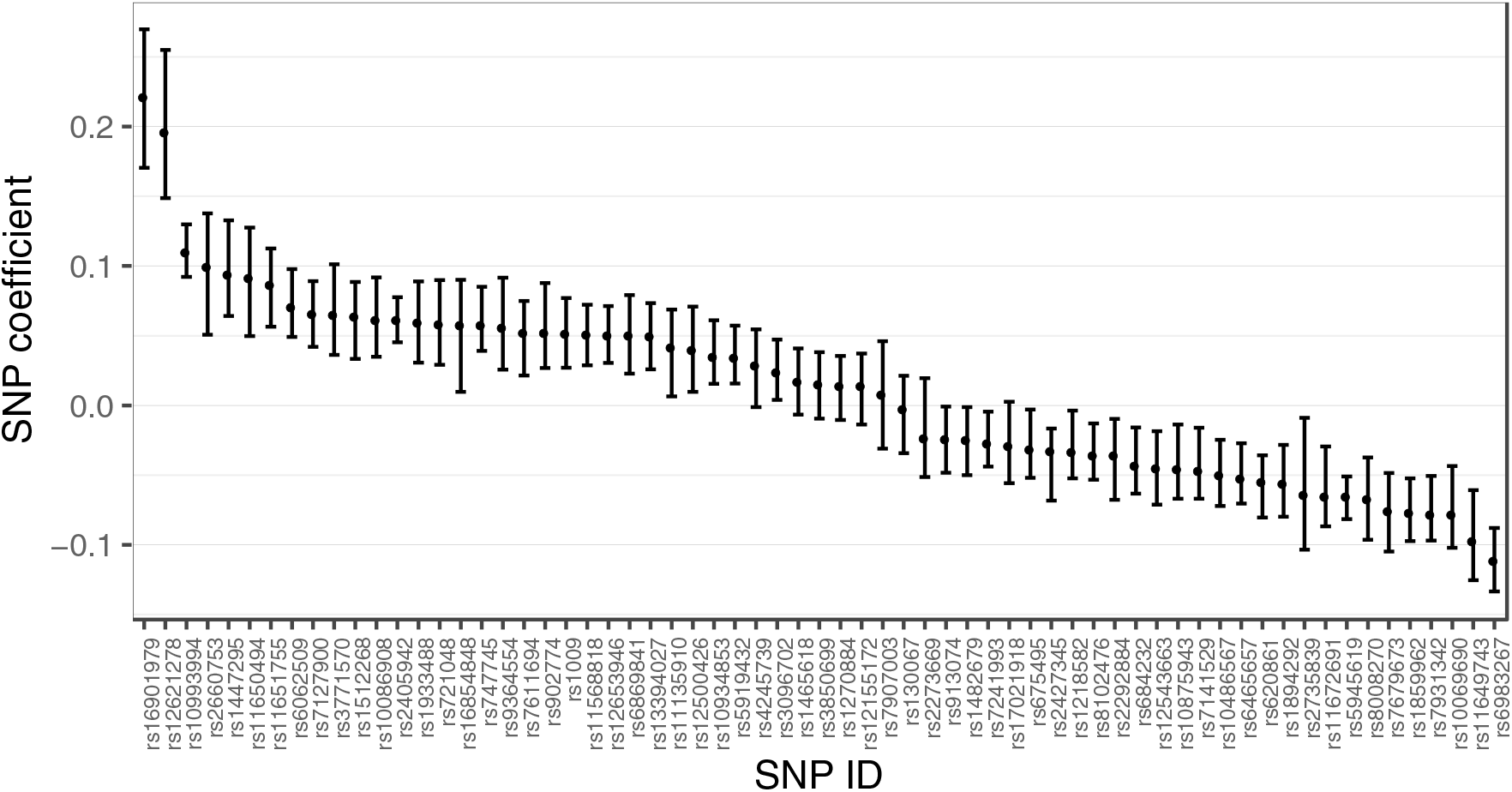
Coefficients of 65 SNPs used in the Established SNP model. Data points represent mean values across 50 iterations of a random sample of the training set using a sample size of 30 thousand total observations. Error bars represent 95% confidence intervals.

### Effect of training set sample size on performance

Box plots of the performance metrics of the Established-SNP and Discovery-SNP models for random samples of the training set are shown in Figure 3 and Figure 4, respectively. The mean values of HR_98/50_, HR_20/50_, HR_98/20_, HR_80/20_, z-score, and beta using a random training sample of 1 thousand total observations in the Established-SNP model were 1.73 [95% CI: 1.69-1.76], 0.71 [0.71-0.73], 2.42 [2.35-2.50], 1.96 [1.92-2.01], 9.92 [9.57-10.28], and 0.45 [0.43-0.47] respectively. The corresponding values using a random training sample of 30 thousand total observations were 2.41 [95% CI: 2.40-2.43], 0.60 [0.60-0.60], 4.04 [4.02-4.07], 2.86 [2.84-2.87], 15.1 [15.04-15.16], and 1.18 [1.17-1.18] respectively.

**Figure 3.**
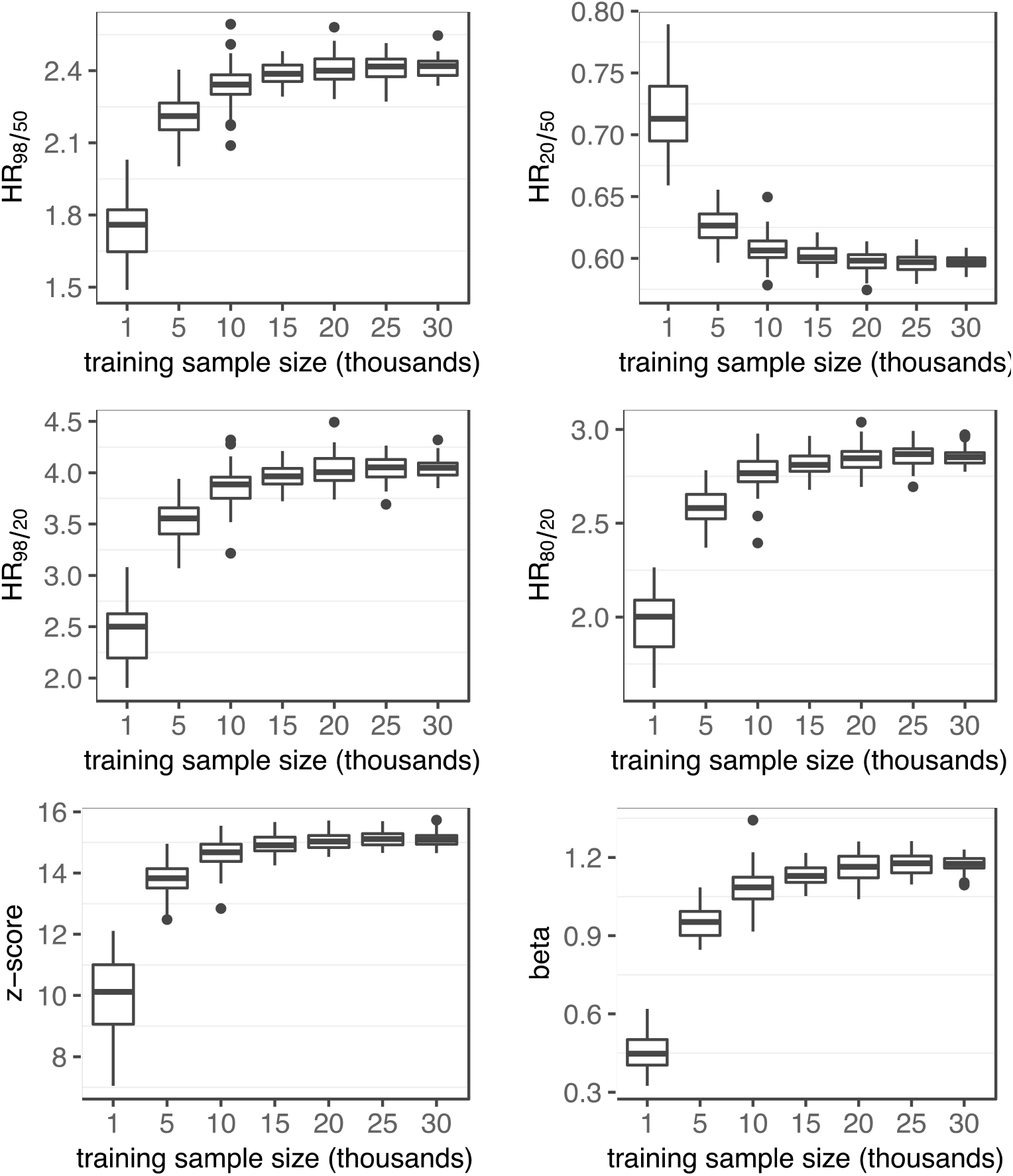
Performance metrics of Established SNP model. Box plots of performance metrics are shown for random samples of the training set using sample sizes of 1, 5, 10, 15, 20, 25, and 30 thousand total observations. Within each box plot, the horizontal line represents the median and the box extends from the 25^th^ to 75^th^ percentile.

**Figure 4.**
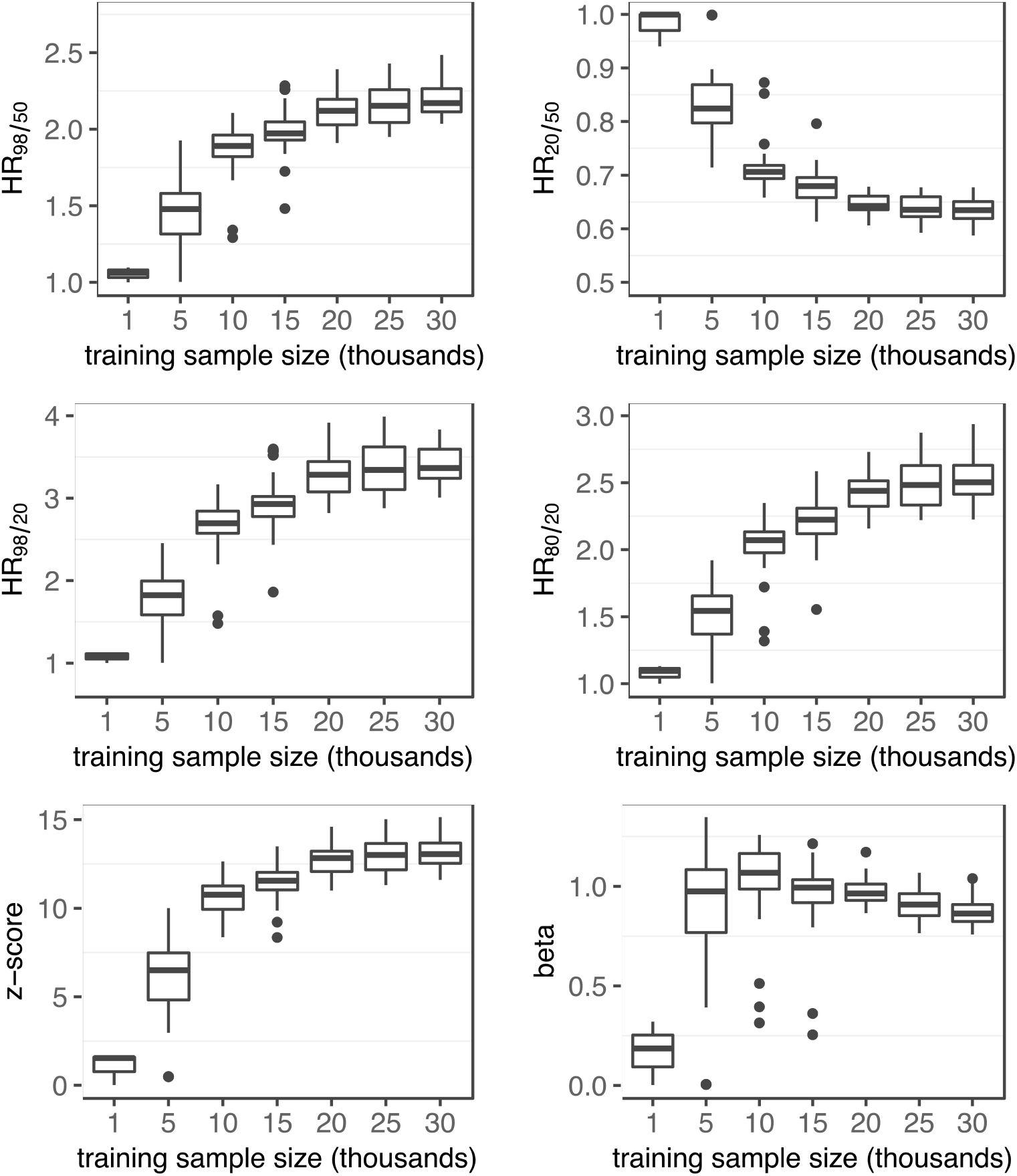
Performance metrics of the Discovery SNP model. Box plots of performance metrics are shown for random samples of the training set using sample sizes of 1, 5, 10, 15, 20, 25, and 30 thousand total observations. Within each box plot, the horizontal line represents the median and the box extends from the 25^th^ to 75^th^ percentile.

The mean values of HR_98/50_, HR_20/50_, HR_98/20_, HR_80/20_, z-score, and beta using a random training sample of 1 thousand total observations in the Discovery-SNP model were 1.05 [0.93-1.18], 0.98 [0.89-1.07], 1.07 [0.91-1.24], 1.08 [0.91-1.24], 1.06 [−1.20-3.31], and 0.17 [−0.23-0.65] respectively. The corresponding performance values using a training sample size of 30 thousand observations were 2.20 [2.16-2.23], 1.60 [1.59-1.62], 3.47 [3.39-3.56], 2.53 [2.49-2.58], 13.19 [12.96-13.41], and 0.87 [0.85-0.89] respectively.

### Effect of testing set sample size on performance

Box plots of the performance metrics of the representative Established-SNP model for random samples of the testing set are shown in Figure 5. The mean values of HR_98/50_, HR_20/50_, HR_98/20_, HR_80/20_, z-score, and beta using a random testing sample of 0.5 thousand total observations in the representative Established-SNP model were 1.78 [1.71-1.85], 0.73 [0.71-0.74], 2.50 [2.33-2.66], 1.99 [1.89-2.09], 3.82 [3.57-4.08], and 0.76 [0.70-0.82] respectively. The corresponding values using a testing sample of 6 thousand observations were: 1.73 [1.72-1.76], 0.73 [0.72-0.73], 2.39 [2.34-2.44], 1.93 [1.90-1.96], 13.07 [12.80-13.32], and 0.74 [0.72-0.76] respectively.

**Figure 5.**
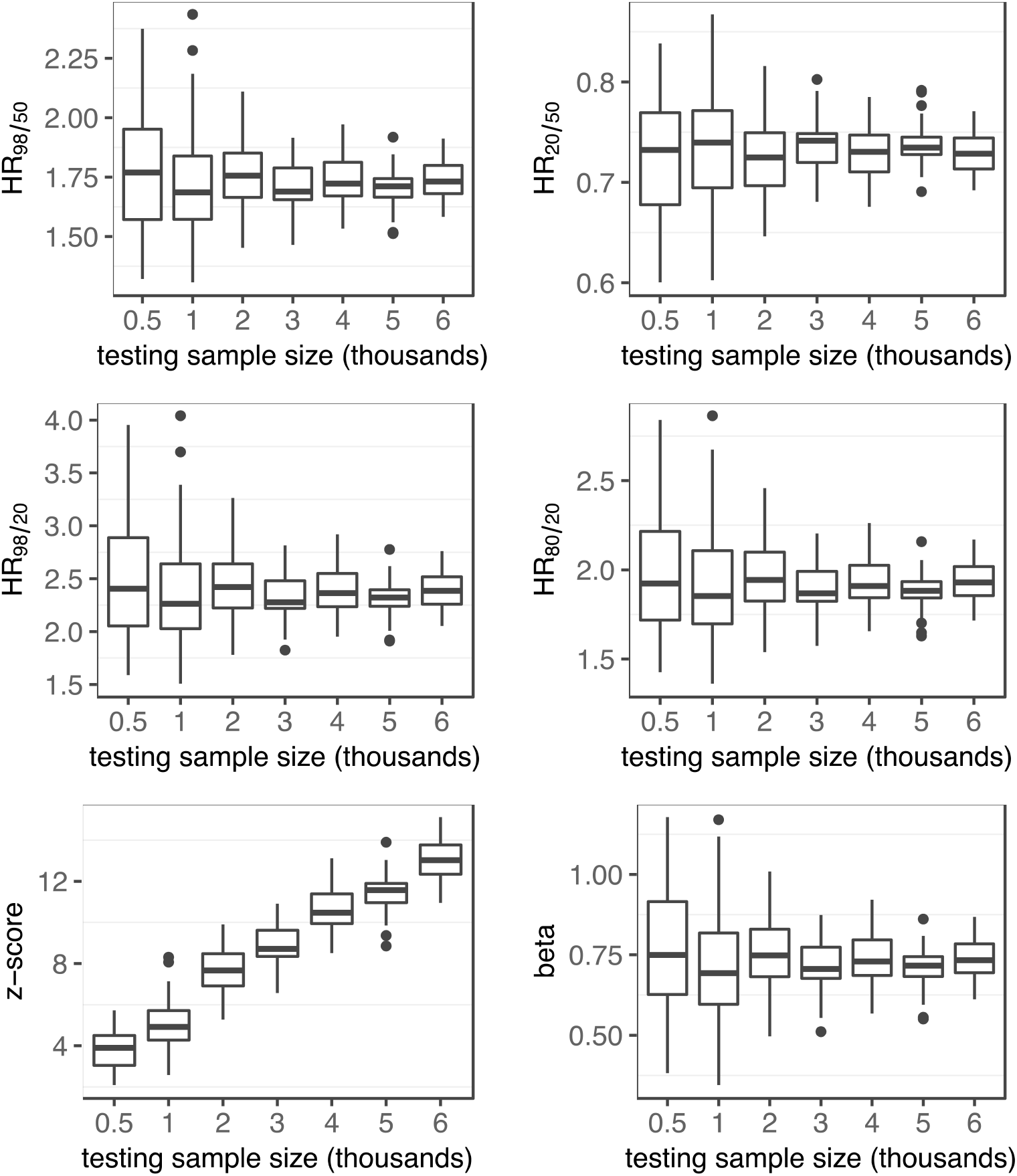
Performance as a function of testing sample size. Box plots of performance metrics of the representative Established SNP model in random samples of the testing set from 0.5 to 6 thousand total observations.

## Discussion

We identified several trends in the effect of training and testing sample size on the performance of PHS models in predicting the age of onset of prostate cancer using SNP genetic variants. When using SNPs that had already been associated with prostate cancer risk, our analysis suggests that very little improvement in performance can be achieved once the training sets becomes larger than 10 to 15 thousand observations. When attempting to discover SNPs, a similar plateau in performance was observed from training sets larger than 20 to 25 thousand observations. Apart from z-scores, the performance metrics of the chosen Cox proportional hazards model did not vary with testing sample size. However, we did observe that the distribution of performance metrics narrows until a testing sample size of 3 to 4 thousand observations, after which the distribution remains relatively stable.

Our results may be used to inform researchers on the approximate number of subjects needed to develop PHS models to predict the age of onset of diseases using SNP counts. A dataset of 20 thousand observations may be the minimum needed to accurately estimate the PHS coefficients of SNPs that have been previously discovered in the setting of a logistic model. Such a dataset would allow for the accurate estimation of SNP coefficients as well as the testing of model performance in an independent holdout set. Based on our results, this number would have to be increased to roughly 30 thousand observations if the researchers intend on discovering the SNPs from scratch using the approach described here.

The PHS model developed by Desikan *et al.*[6] to predict age-associated risk of Alzheimer’s disease used a training set with roughly 55,000 individuals. A similarly structured model developed by Seibert *et al.*[7] to guide screening for aggressive prostate cancer was developed with roughly 31,000 men. Studies such as these require large investments in time, money, and resources in order to acquire the genetic data needed for the analysis. The results of our analysis help elucidate that the minimum sample size needed to translate this technology to other diseases and processes may be lower than what has been used so far in previous studies. This seems to be particularly true if the researchers use SNPs that have already been discovered and validated as associated with the process of interest.

The results of this study must be considered in the context of its limitations. The list of Established-SNPs was previously selected from a larger dataset that included the sample patients used in the test set in the present study. As such, there is leakage of information from the test set to the development of the Established-SNP model. Therefore, the performance metrics of the Established-SNP model should not be directly compared to those of the Discovery-SNP model, as the values of the former may be inflated.

In addition, we have chosen to focus on only two of countless possible model development schemes. The role of sample size in other development strategies—such as regularized Cox proportional models, parametric survival functions, or random survival forests—is yet to be explored. Finally, the analysis is limited to prostate cancer and to the SNPs on the iCOGS array. Future work will include SNPs imputed from 1000 Genomes[13]. Such an analysis was not performed for this first study to limit computation time for bootstrap analyses and to avoid uncertainty due to imputation.

In conclusion, we have studied the effect of sample size on the performance of PHS models to study the association between SNPs and the age at onset of prostate cancer. We have determined that models require roughly 20 to 30 thousand samples before their performance would not be improved greatly by expansion of the training set. Using SNPs that have already been established in the literature may help reduce the number of training samples required to reach this performance plateau by almost 10 thousand samples.

## Materials and Methods

### Training and testing set

As previously described[7], we obtained genotype and age data from 21 studies included in the Prostate Cancer Association Group to Investigate Cancer Associated Alterations in the Genome (PRACTICAL) consortium. We analyzed data from 40,861 men consisting of 20,551 individuals with prostate cancer and 20,310 individuals without. For analysis, the age for each man was recorded as either their age at prostate cancer diagnosis (cases) or at interview (controls). Genotype data for 201,590 SNPs were also available for analysis. The genotype data had been assayed using a custom iCOGS chip (Illumina, San Diego, CA) the details for which are elaborated elsewhere[10]. The sample was split into training (34,444 men) and testing (6,417 men) sets. The testing set was selected using men who were enrolled in the Prostate testing for cancer and Treatment (ProtecT[11]) trial. ProtecT (ClinicalTrials.gov: NCT02044172) is a large, multicenter trial within the United Kingdom which aims to investigate the effectiveness of treatments for localized prostate cancer. The ProtecT study group was chosen for testing as it represented a well-characterized group of individuals that had been used for measuring testing performance for our earlier work. The Data Availability Statement describing how readers can gain access to the PRACTICAL dataset is provided in the Supplementary Information.

### Established-SNP model

A list of 65 SNPs[12] was chosen to represent those on the iCOGS array that had been published as associated with prostate cancer. The coefficients of the SNPs within the Established-SNP model were then estimated using the “coxphfit” function in MATLAB (Mathworks, Natwick, MA). Prior to parameter estimation, missing SNP data were replaced by mean imputation. It should be noted that the 65 SNPs used were discovered, in large part, using the data presently defined as the test set. The effect allele for all 65 SNPs was defined as “A” to simplify analysis.

### Discovery-SNP model

SNPs with call rates less than 95% were removed from the selection process. For every SNP, a trend test was used to check for associations between SNP count and the binary classification of individuals with or without prostate cancer. The SNP selection pool was then reduced to those whose trend test p-value was less 1×10^−6^. In order of increasing p-value, each SNP was tested in a multiple logistic regression model for association with the binary classification of men as with or without prostate cancer, after adjusting for age, six principal components based upon genetic ancestry, and previously selected SNPs. If the p-value of the coefficient of the tested SNP was less than 1×10^−6^, it was selected for the final Cox proportional hazard model estimation. The coefficients of the selected SNP pool within the Discovery-SNP model were estimated as previously described[7].

### Polygenic Hazard Score (PHS)

The polygenic hazard score (PHS) for each of the Established-SNP and Discovery-SNP models was calculated as the linear product of the coefficients of the SNPs used in the model and the corresponding patient genotype counts[6,7].

### PHS performance metrics

Several performance metrics for PHS models were investigated, and are described in Table 1. In each case, the PHS for each test subject was calculated as the dot product of SNP coefficients, either Established or Discovery, and SNP counts. A Cox proportional hazards model was then fit using PHS as the sole predictor of age in the test set. The z-score and beta of this Cox proportional hazards model relate to how well PHS was associated with age within the test set. The hazard ratios were calculated as the exponential of the differences in predicted log-relative hazards of different groups within the test set. The groups were defined using centile cut-points for those controls within the training set whose age was less than 70 years. This list of performance metrics expands on those (z-score and HR_98/50_) that were used in our earlier work[7].

**Table 1.** Performance metrics used in the evaluation of polygenic hazard scores.

| Performance metric | Description |
| --- | --- |
| $HR_{98/50}$ | Hazard ratio of the top 2% to the average (30 – 70%) in the test set |
| $HR_{20/50}$ | Hazard ratio of the bottom 20% to the average (30 – 70%) in the test set |
| $HR_{98/20}$ | Hazard ratio of the top 2% to the bottom 20% in the test set |
| $HR_{80/20}$ | Hazard ratio of the top 20% to the bottom 20% in the test set. |
| z-score | z-score of Cox proportional hazards model using PHS as a sole predictor of age in the test set |
| beta | coefficient of PHS in a Cox proportional hazards model using PHS as a sole predictor of age in the test set. |

### Random sampling of training set

Random sampling of the training set was performed with replacement while ensuring equal proportions of men with and without prostate cancer. The training set was randomly sampled to include 1, 5, 10, 15, 20, 25, and 30 thousand total observations. Performance of the Established and Discovery-SNP models using random samples of the training data was measured in the entire test set.

### Random sampling of the testing set

Random sampling of the testing set was performed with replacement while ensuring equal proportion of men with and without prostate cancer. The testing set was randomly sampled to include 0.5, 1, 2, 3, 4, 5 and 6 thousand total observations. Performance in the randomly sampled testing sets was performed using a representative Established-SNP model. The representative model was chosen as that whose parameters were estimated using a training sample size of 30 thousand total observations, and whose performance metrics were the shortest Euclidean distance to the average performance across all Established-SNP models using a training sample size of 30 thousand.

## Supporting Information Legends

Supporting Information 1. Data Availability Statement details how readers can obtain the data from the PRACTICAL (Prostate Cancer Association Group to Investigate Cancer Associated Alterations in the Genome) consortium. The document also contains the additional authorship, affiliation, and funding sources for the PRACTICAL consortium.

